# Acute IFN-γ responses drive sensory neuron injury and chronic pain after chikungunya virus infection

**DOI:** 10.64898/2026.02.02.702518

**Authors:** Lilian C. Colodeti, Thamires B. P. Gonçalves, Hannah P. Mota-Araujo, Suzana M. B. Araujo, Greice N. Pires, Bruno V. M. Santiago, Thiago S. Bacelar, Fabricia L. Fontes-Dantas, Emily C.C. Rennô, Mariana O. L. da Silva, Luiz Eduardo B. Savio, Pedro S. Ciriaco, Tiago B. Taboada, Ana B. O. Barbosa, Fabio M. Gomes, Ana M. B. Martinez, Luciana J. Costa, Iranaia Assunção-Miranda, Nivaldo R. Villela, José L. Proença-Modena, Henrique R. Mendonça, Soniza Alves-Leon, Giselle F. Passos, William M. de Souza, Robson da Costa, Claudia P. Figueiredo

## Abstract

Chikungunya virus (CHIKV) is a mosquito-borne alphavirus that causes acute and chronic musculoskeletal disease characterized by often debilitating pain; however, the mechanisms associated with the pain remain understudied. Here, we used an experimental mouse model and patients to investigate the role of interferon-γ (IFN-γ) in the pain during chronic inflammation after CHIKV infection. Our data show that CHIKV induces sustained joint inflammation, cartilage catabolism, and sensory neuron injury signatures, including *Atf3* upregulation, but lacks detectable viral dissemination to dorsal root ganglia. We found that IFN-γ expression was higher in CHIKV-infected joints, and direct IFN-γ administration recapitulates mechanical hypersensitivity and neuronal stress independently of joint degeneration. Genetic or pharmacological disruption of IFN-γ signalling prevented CHIKV-induced pain and neuronal stress without altering viral burden, demonstrating that IFN-γ is required for both sensory dysfunction and joint pathology. By stratifying CHIKV-infected mice based on long-term nociceptive profiles, we show that persistent-pain subgroup characterized by elevated acute-phase systemic IFN-γ and chronic peripheral neuropathic features. These findings were confirmed in human patients, where acute-phase IFN-γ levels distinguished those who developed chronic arthralgia from those who recovered. Overall, our findings demonstrate that persistent post-CHIKV pain is driven by an IFN-γ-mediated neuroimmune mechanism.

## Introduction

Chikungunya virus (CHIKV) is a mosquito-borne alphavirus that causes debilitating acute and chronic disease, and some cases can lead to neurological manifestations and death^1-6^. Persistent pain represents one of the most disabling sequelae of chikungunya infection and a major driver of its long-term socioeconomic burden^7,8^. Clinically, polyarthralgia and polyarthritis can persist for weeks (subacute phase), or from months to years (chronic phase)^1^. Currently, CHIKV causes approximately 17.7 million annual cases across 104 countries, and nearly 2.8 billion people live in areas at risk of infection^7^. In 2025, autochthonous CHIKV cases and outbreaks were reported in France, Italy, China, and the United States of America (USA), demonstrating continued geographic expansion^9-13^. Consequently, chikungunya has generated substantial public health and socioeconomic burden, with estimated global costs of $2.8 billion (direct) and $47.1 billion (indirect) between 2011 and 2020^14^. Between 2023 and 2025, Ixchiq (live attenuated) and Vimkunya (virus-like particle) vaccines were licensed, but the deployment of immunization in endemic countries remains under debate^15^.

At present, the chronic phase, characterized by often debilitating pain, is the most disabling consequence of CHIKV infection; however, its underlying mechanisms remain poorly understood. Some studies suggest that chronic phases are associated with higher levels of IL-6 and IL-17, which may disrupt the RANKL/OPG ratio and promote bone erosion^16,17^. Interestingly, higher levels of interferon-γ (IFN-γ) are observed during acute CHIKV infection in both clinical studies and experimental studies^18-21^. Notably, IFN-γ has been demonstrated to be a biological driver of neuropathic pain, a highly debilitating pain condition that commonly occurs after nerve damage^22,23^. However, the role of IFN-γ in CHIKV-associated pain remains elusive. In this study, we combine an experimental mouse model and a patient cohort to investigate the role of IFN-γ in pain associated with CHIKV infection.

## Results

### CHIKV infection triggers joint pathology and sensory neuron stress

To dissect the pathophysiology of CHIKV-induced pain, we developed an experimental mouse model by intra-articular inoculation of CHIKV into the knee joint to recapitulate viral tropism observed in patients. We confirmed CHIKV replication in the ipsilateral knees up to 3 days post-infection (dpi), and CHIKV RNA was detectable up to 28 dpi (**Fig. 1a**). Immunolabeling of double-stranded RNA (dsRNA) indicated the persistence of CHIKV within articular cartilage and synovial tissue up to 28 dpi, whereas mock-infected controls showed no signal (**Fig. 1b**). CHIKV-infected mice exhibited significantly reduced ipsilateral paw withdrawal thresholds starting at 10 dpi and persisting through 28 dpi compared with mock-infected controls (**Fig. 1c**). Conversely, cold sensitivity did not differ significantly between groups, and no sex-dependent difference was observed in either assay (**Fig. 1d and Extended Data Fig. 1a-d**).

**Fig. 1.**
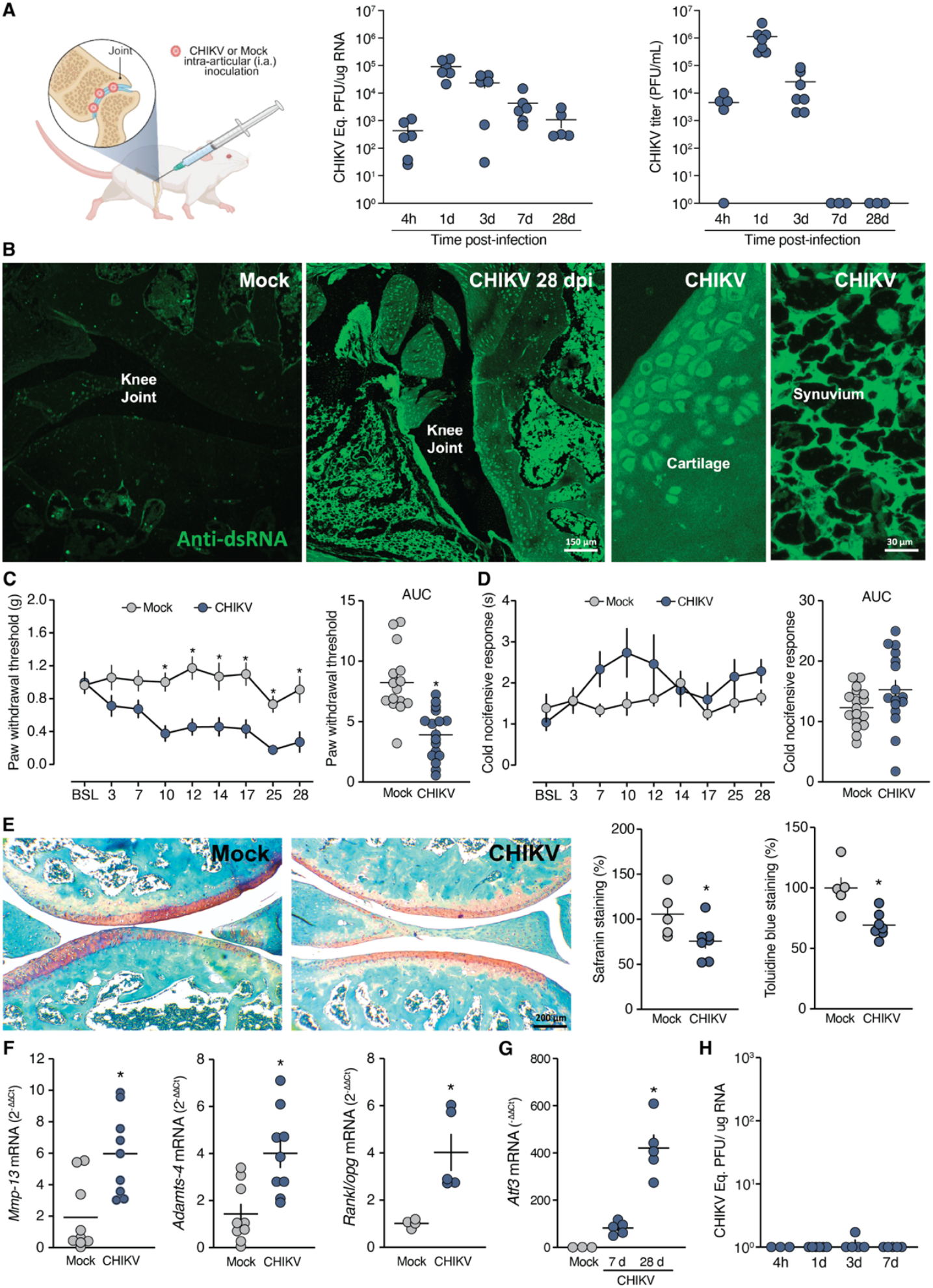
CHIKV infection induces joint pathology and transient neuronal stress without persistent viral RNA in sensory ganglia. **a**, Intra-articular CHIKV inoculation (10^6^ PFU) and viral burden in ipsilateral joints measured by qPCR and plaque assay over time. **b**, dsRNA immunostaining in joint tissues at 28 dpi. **c**, Mechanical hypersensitivity (von Frey) and **d**, cold nocifensive responses (acetone test) following infection, with corresponding AUC. **e**, Representative Safranin O staining of knee joint sections at 28 dpi with corresponding quantification, and quantification of toluidine blue staining. **f**, Expression of joint catabolic markers (*Mmp13, Adamts4, Rankl/opg* ratio) at 28 dpi. **g**, *atf3* mRNA expression. **h**, Viral RNA levels in dorsal root ganglia (DRG). Data are the mean and standard error. In **c-d**, 15–17 mice per group, two-way ANOVA, AUC analyzed by two-tailed unpaired *t* test; **e-f**, 4–9 mice per group, two-tailed unpaired t test. **g**, 3–5 mice per group, Kruskal-Wallis test. *P < 0.05.

Histological analysis revealed that CHIKV infection led to marked synovial hyperplasia, dense inflammatory infiltrates, and disorganization of knee tissue compared with mock controls at 7 and 28 dpi (**Extended Data Fig. 2a**). We further observed a significant loss of proteoglycans in CHIKV-infected mice, as indicated by reduced safranin-O and toluidine blue staining at 28 dpi (**Fig. 1e and Extended Data Fig. 2b**). Consistent with these findings, expression of joint catabolic markers (*Mmp13, Adamts4*, and the *Rankl/Opg* mRNA ratio) was significantly increased in CHIKV-infected knees at 28 dpi (**Fig. 1f**). To determine if the observed mechanical allodynia involved a neuropathic component, we tested sensitivity to gabapentin (GBP)^24^. GBP robustly attenuated mechanical allodynia and reduced overall hypersensitivity (**Extended Data Fig. 2c)**. Moreover, activating transcription factor 3 (*Atf3*) expression was significantly upregulated in the lumbar dorsal root ganglia (DRG) at 28 dpi in CHIKV-infected mice (**Fig. 1g**), indicating sustained neuronal stress. CHIKV RNA was undetectable in lumbar DRGs up to 7 dpi, ruling out direct viral infection of the ganglia (**Fig. 1h**). These results indicate that CHIKV infection induces indirect sensory neuron injury and supports a neuropathic contribution to the persistent pain state.

### IFN-γ and T cells drive joint inflammation in CHIKV-induced arthritis

We next characterized the expression of pro-inflammatory mediators and the corresponding cellular immune response. qPCR analysis of the knee tissue showed transient induction of *il-6* and *tnf-α* mRNA, peaking at 3 and 7 dpi, respectively. Type I interferons were induced early, with *ifn-α* and *ifn-β* expression peaking at 1 dpi before returning to baseline. Notably, *ifn-γ* expression was strongly and persistently elevated, peaking at 7 dpi and remaining above mock-infected control levels through 28 dpi (**Fig. 2a**). To identify the immune cell populations driving this inflammation, we profiled synovial fluid and peripheral blood via flow cytometry. Joint fluid analysis revealed a pronounced influx of CD4^+^ and CD8^+^ T cells at 7 dpi, which largely resolved by 28 dpi. Myeloid cells (CD11b^+^CD45^+^) displayed a similar kinetic profile, peaking in the joint at 7 dpi and returning towards baseline at 28 dpi. This joint infiltration occurred concomitantly with a significant reduction in circulating CD11b^+^CD45^+^ cells at 7 dpi, consistent with their recruitment from blood into the inflamed joint (**Fig. 2b-d and Extended Data Fig. 3a**). We further sought to identify the specific lymphocyte populations responsible for IFN-γ production within the joint. IFN-γ expression by NK cells remained lower (<2% IFN-γ^+^ cells) at 7 and 28 dpi, whereas IFN-γ^+^ CD8^+^ T cells were markedly increased at 7 dpi (**Fig. 2c and Extended Data Fig. 3b**). IFN-γ^+^ CD4^+^ T cells also increased at 7 dpi, albeit to a lesser extent than their CD8^+^ counterparts (**Extended Data Fig. 3b**). These data establish CD8^+^ T cells as the predominant IFN-γ-producing lymphocyte population in the joint during the peak of inflammation.

**Figure 2.**
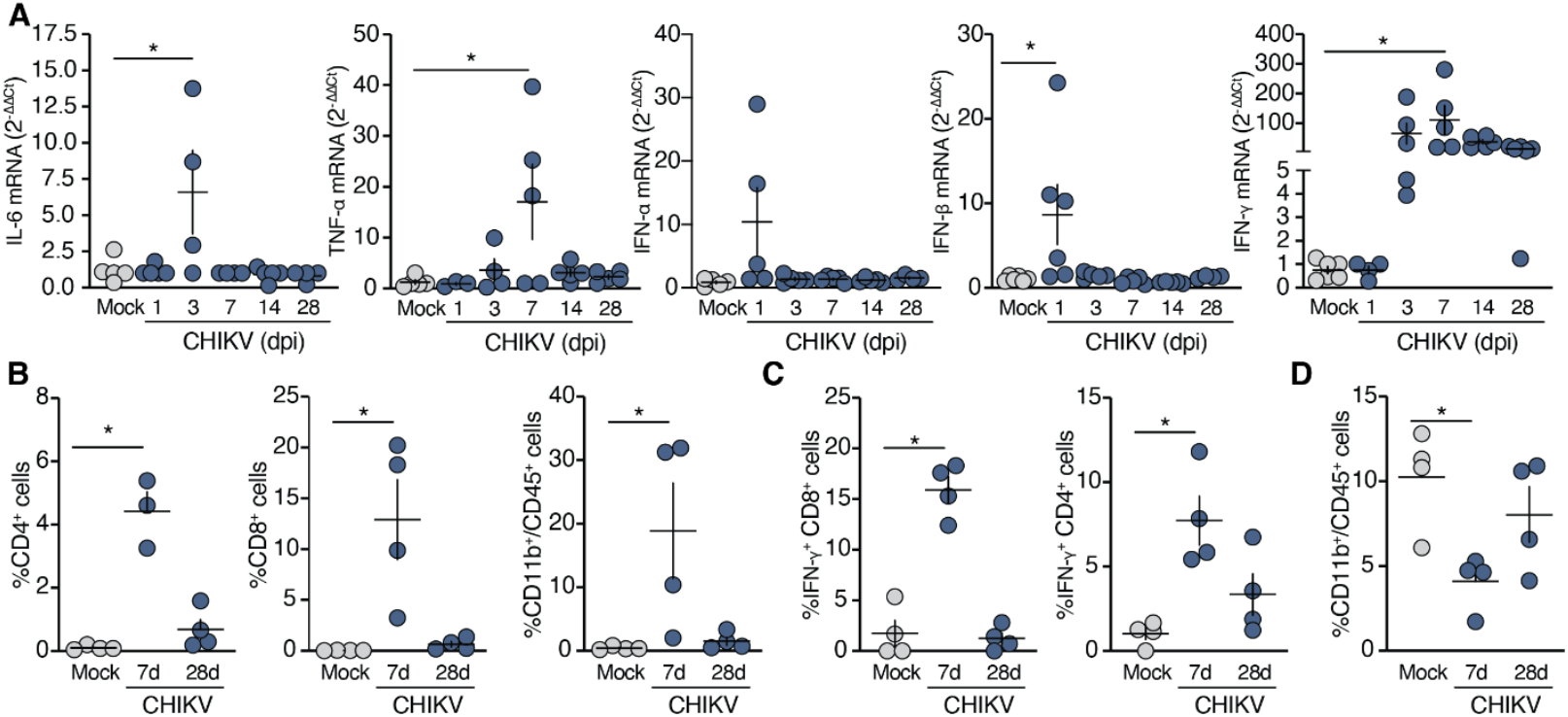
IFN-γ response and T cell recruitment in joint inflammation after CHIKV infection. **a**, qPCR analysis of pro-inflammatory mediators in ipsilateral knee joints following intra-articular CHIKV infection. **b**, Quantification of leukocyte populations in synovial fluid at 7 and 28 dpi. **c**, Frequencies of IFN-γ–producing CD8^+^ and CD4^+^ T cells in synovial fluid. **d**, Quantification of leukocyte populations in blood at 7 and 28 dpi. Data are the mean and standard error. In **a**, 3–6 mice per group, one-way ANOVA followed by Tukey’s multiple-comparisons test; **b**,**c** and **d**, 3–4 mice per group, Kruskal-Wallis test. *P < 0.05.

### IFN-γ drives neuronal stress and pain in CHIKV infection

To assess whether IFN-γ triggers joint pain hypersensitivity, we administered intra-articular recombinant IFN-γ to uninfected mice. We observed that repeated dosing (300 or 600 U/site every 2 days for 7 days) induced rapid and sustained mechanical allodynia in the von Frey test (**Fig. 3a**). Thus, we used a lower dose (300 U/site) for subsequent experiments. Despite this robust pain phenotype, the expression of *Adamts4* and *Mmp13* remained unchanged relative to vehicle-treated controls, indicating that IFN-γ causes pain independently of structural joint tissue remodelling (**Fig. 3b**). Consistently, intra-articular IFN-γ significantly increased *Atf3* expression in lumbar DRGs, supporting a direct capacity of IFN-γ to induce sensory neuron stress and pain without joint degeneration (**Fig. 3c**). We next asked whether IFN-γ can act on sensory neurons to elicit activation and stress signalling. To address this, primary mouse DRG neurons were cultured and exposed to IFN-γ, and ERK phosphorylation (p-ERK) and ATF3 induction were assessed as a readout of neuronal activation and stress. We found that IFN-γ treatment led to a significant increase in p-ERK^+^ neurons, similar to that observed with capsaicin, a positive control (**Fig. 3d**). IFN-γ exposure also increased the proportion of ATF3^+^ neurons, as demonstrated by immunostaining and qPCR analysis (**Fig. 3e**). Together, these results indicate that IFN-γ directly activates DRG neurons and induces a transcriptional stress response.

**Fig. 3.**
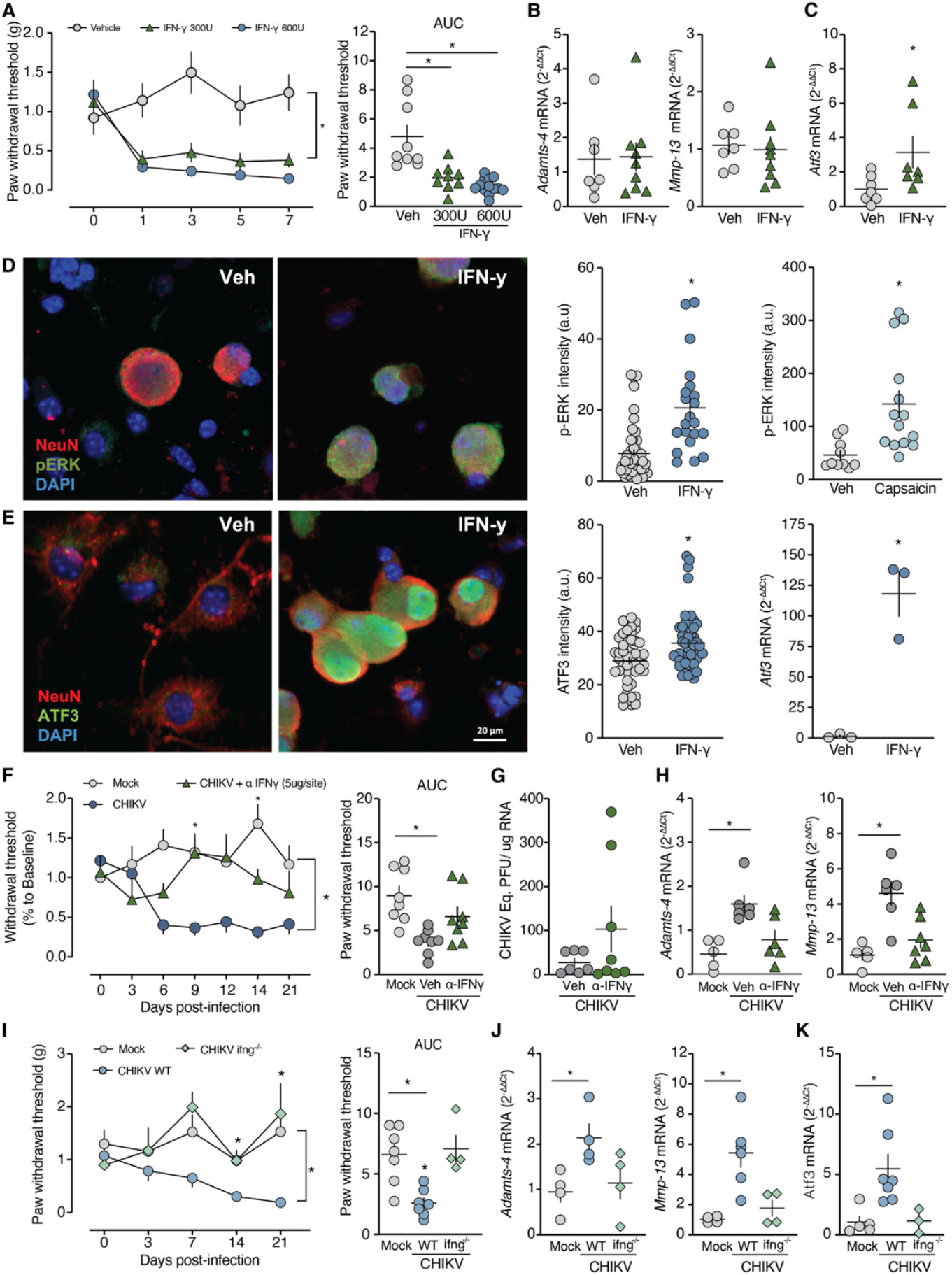
IFN-γ signaling drives CHIKV-induced pain and neuronal stress. **a**, Intra-articular administration of recombinant IFN-γ (300 or 600 U per site; days 0, 2, and 4) induced mechanical withdrawal thresholds shown as time course and AUC. **b**, Joint *Adamts4* and *Mmp13* mRNA expression following IFN-γ administration. **c**, DRG Atf3 mRNA levels following IFN-γ administration. **d**, p-ERK immunoreactivity in cultured NeuN^+^ DRG neurons after IFN-γ exposure; capsaicin served as a positive control. **e**, ATF3 immunoreactivity and *atf3* mRNA levels in cultured DRG neurons following IFN-γ exposure. **f**, Mechanical withdrawal thresholds following treatment with an IFN-γR1–blocking antibody (anti-CD119) during CHIKV infection. **g**, Joint CHIKV RNA levels after anti-CD119 treatment. **h**, Joint *Adamts4* and *Mmp13* expression after anti-CD119 treatment. **i**, Mechanical withdrawal thresholds in wild-type (WT) and ifng^-^/^-^ mice following CHIKV infection (time course and AUC). **j**, Joint *Adamts4* and *Mmp13* expression in WT and ifng^-^/^-^ mice following CHIKV infection. **k**, DRG *atf3* mRNA levels in WT and ifng^-^/^-^ mice. Data are the mean and standard error. In **a, f** and **i**, 8–14 mice per group, two-way ANOVA followed by Tukey’s multiple-comparisons test; AUC analyzed by two-tailed unpaired *t* test; **h, j** and **k**, 5–9 mice per group, one-way ANOVA followed by Tukey’s multiple-comparisons test; **b, c** and **g**, 5–7 mice per group, two-tailed unpaired *t* test; **d** and **e**, three independent neuronal cultures (two coverslips per culture), two-tailed unpaired *t* test. *P < 0.05.

We next tested whether blocking IFN-γ signalling attenuates pain development after CHIKV infection and examined the impact on nociceptive behaviour and DRG stress responses. We observed that intra-articular administration of the anti-CD119 monoclonal antibody (IFN-γ receptor 1) significantly attenuated CHIKV-induced mechanical allodynia compared with non-treated CHIKV-infected mice, while viral RNA levels in the joint remained unchanged at 21 dpi (**Fig. 3f-g**). Anti-CD119 treatment also reduced CHIKV-induced upregulation of *Adamts4* and *Mmp13* (**Fig. 3h**). Consistently, *Ifng*^-/-^ mice (lacking expression of *ifn-*γ) were similarly protected, maintaining baseline withdrawal thresholds through 21 dpi, with no change in joint viral RNA (**Fig. I and Extended Data Fig. 4a-b**). Notably, *the absence of Ifng* abolished the induction of A*damts4* and *Mmp13* in the joint and *Atf3* in the DRG (**Fig. 3j-k**), indicating that IFN-γ signalling is required for both joint catabolic responses and neuronal stress associated with CHIKV infection. These findings identify IFN-γ signalling as a key contributor to CHIKV-induced pain, independent of viral clearance.

### Acute systemic IFN-γ responses predict persistent pain after CHIKV infection

To determine whether the acute and chronic phases of CHIKV infection are recapitulated in our intra-articular CHIKV infection model and to probe the contribution of IFN-γ to chronic pain, we extended our analyses to 77 dpi. Mechanical sensitivity assessed by the von Frey test revealed that CHIKV-infected mice had significantly reduced ipsilateral paw withdrawal thresholds from 14 to 42 dpi compared with mock-infected controls (**Fig. 4a**). We next stratified CHIKV-infected mice based on their mechanical sensitivity profiles. Animals with a ≥30% reduction from baseline (i.e., withdrawal thresholds ≤70% of baseline) were categorized as the persistent group, whereas those with a <30% reduction (>70% of baseline) were classified as the recovered group (**Fig. 4b**). We found that the persistent group (12 of 24 animals) maintained significantly lower withdrawal thresholds through 77 dpi, indicating long-term pain, whereas recovered animals returned to baseline sensitivity by 49 dpi. During the chronic phase (42–77 dpi), the persistent group exhibited a significantly lower withdrawal threshold than mock-infected controls and recovered animals (**Fig. 4c**).

**Fig. 4.**
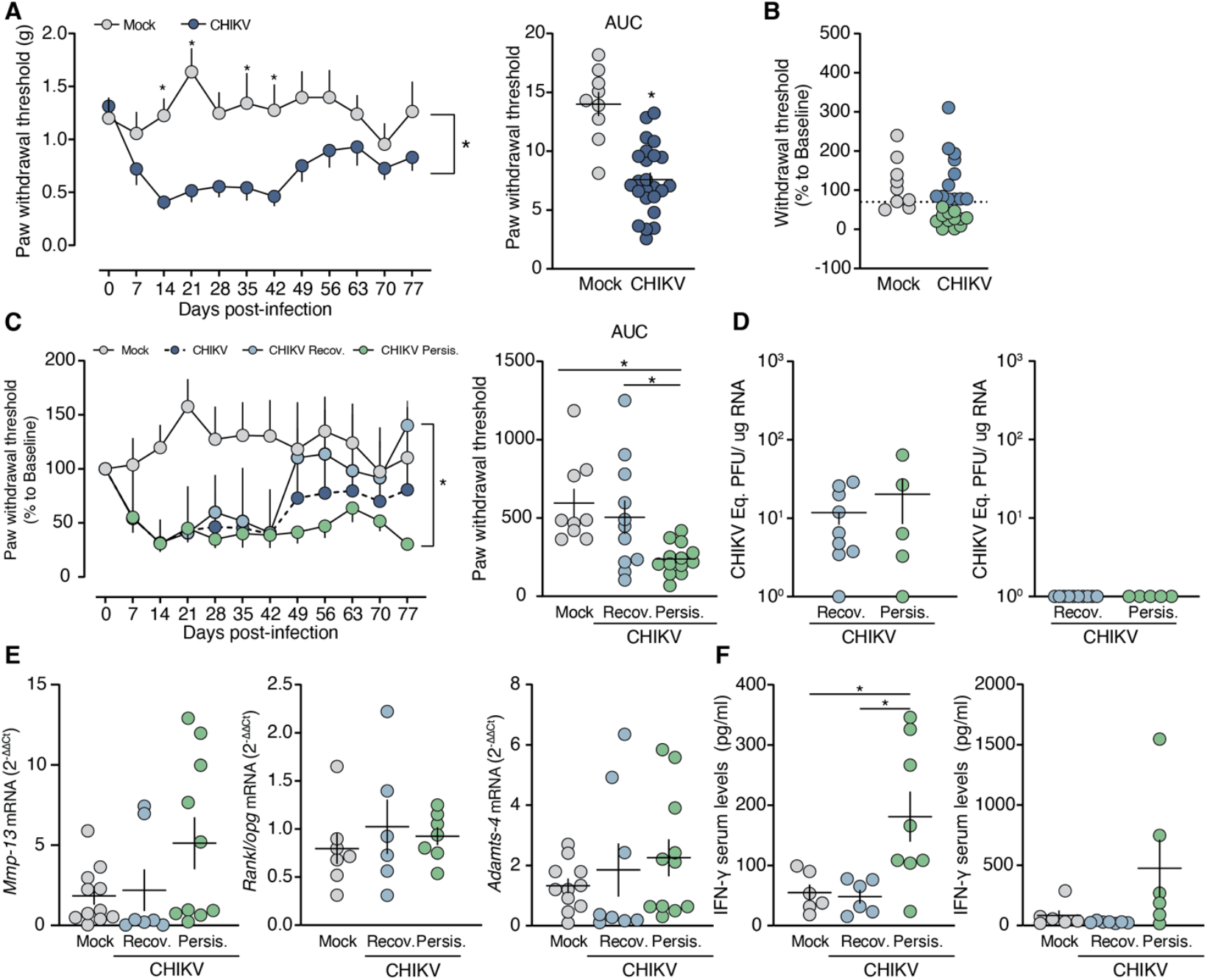
Persistent pain after CHIKV infection is associated with elevated acute IFN-γ responses but not sustained viral persistence or joint catabolism. **a**, Time course of paw withdrawal thresholds following CHIKV infection (0–77 dpi), shown with corresponding AUC. **b-c**, Stratification of CHIKV-infected mice at 77 dpi based on mechanical sensitivity into recovered (CHIKV-Recov.) and persistent (CHIKV-Persis.) groups, with longitudinal analysis and late-phase AUC. **d**, CHIKV RNA levels in knee tissue and dorsal root ganglia (DRG) at 77 dpi. **e**, Expression of joint catabolic genes (*Mmp13, Adamts4*) and *Rankl/opg* mRNA ratio at 77 dpi. **f**, Serum IFN-γ concentrations during acute (7 dpi) and chronic phases (77 dpi). Data are the mean and standard error. In **a** and **b**, 9–24 mice per group, two-way ANOVA followed by Tukey’s multiple-comparisons test; AUC analyzed by two-tailed unpaired *t* test; **c**, 9–13 mice per group, two-way ANOVA followed by Tukey’s multiple-comparisons test; late-phase AUC analyzed by one-way ANOVA followed by Tukey’s multiple-comparisons test; **d** and **e**, 5–11 mice per group, two-tailed unpaired *t* test; **f**, 6–7 mice per group, one-way ANOVA followed by Tukey’s multiple-comparisons test. *P < 0.05.

To assess whether persistent pain was associated with ongoing viral replication in peripheral tissues, we quantified CHIKV RNA in the ipsilateral knee joints and lumbar DRGs at 77 dpi. CHIKV RNA levels were comparable between mice with persistent and recovered phenotypes (**Fig. 4d**). Similarly, expression of cartilage- and bone-catabolic genes *Mmp13* and *Adamts4*, and the *Rankl*/*Opg* mRNA ratio, were comparable across groups (**Fig. 4e**). These findings demonstrate that persistent pain following CHIKV infection occurs independently of viral persistence or ongoing structural joint degeneration. Then, we asked whether differential IFN-γ responses might underlie the distinct pain outcomes observed in CHIKV-infected mice. Serum IFN-γ concentrations were significantly elevated in the persistent group during the acute phase compared with recovered and mock controls, whereas no differences were observed during the chronic phase (**Fig. 4f**). These data indicate that increased systemic IFN-γ responses during the acute phase of CHIKV infection are associated with long-term pain in mice.

### Persistent CHIKV pain associates with chronic peripheral neuropathic alterations

We next asked whether persistent pain after CHIKV infection is accompanied by molecular and morphological changes in peripheral sensory structures. We observed higher cleaved caspase-3 immunoreactivity in DRG neurons from animals with persistent pain compared to the recovered group (**Fig. 5a**). In contrast, *Atf3* mRNA levels in the DRG did not differ among groups at 77 dpi (**Extended Data Fig. 5a**). To determine whether DRG caspase-3 activation translates into chronic morphological alterations, we examine the sciatic nerve innervating the affected joint. Quantification of myelinated fiber density in semi-thin sciatic nerve cross-sections revealed no significant differences among groups (**Fig. 5b and Extended Data Fig. 5b**). Analysis of myelin morphology revealed discrete alterations, including vacuolization and myelin splitting, in the persistent pain group compared with the mock control (**Fig. 5b and Extended Data Fig. 5c**). Consistently, g-ratio distribution analysis revealed a significant reduction of fibres within the optimal 0.56–0.65 range in the CHIKV-persistent pain group compared with recovered group (**Fig. 5b**). Immunofluorescence analysis of the sciatic nerve revealed a reduction of neurofilament 200 signal intensity in animals with persistent pain compared with mock animals (**Extended Data Fig. 5d**).

**Fig. 5.**
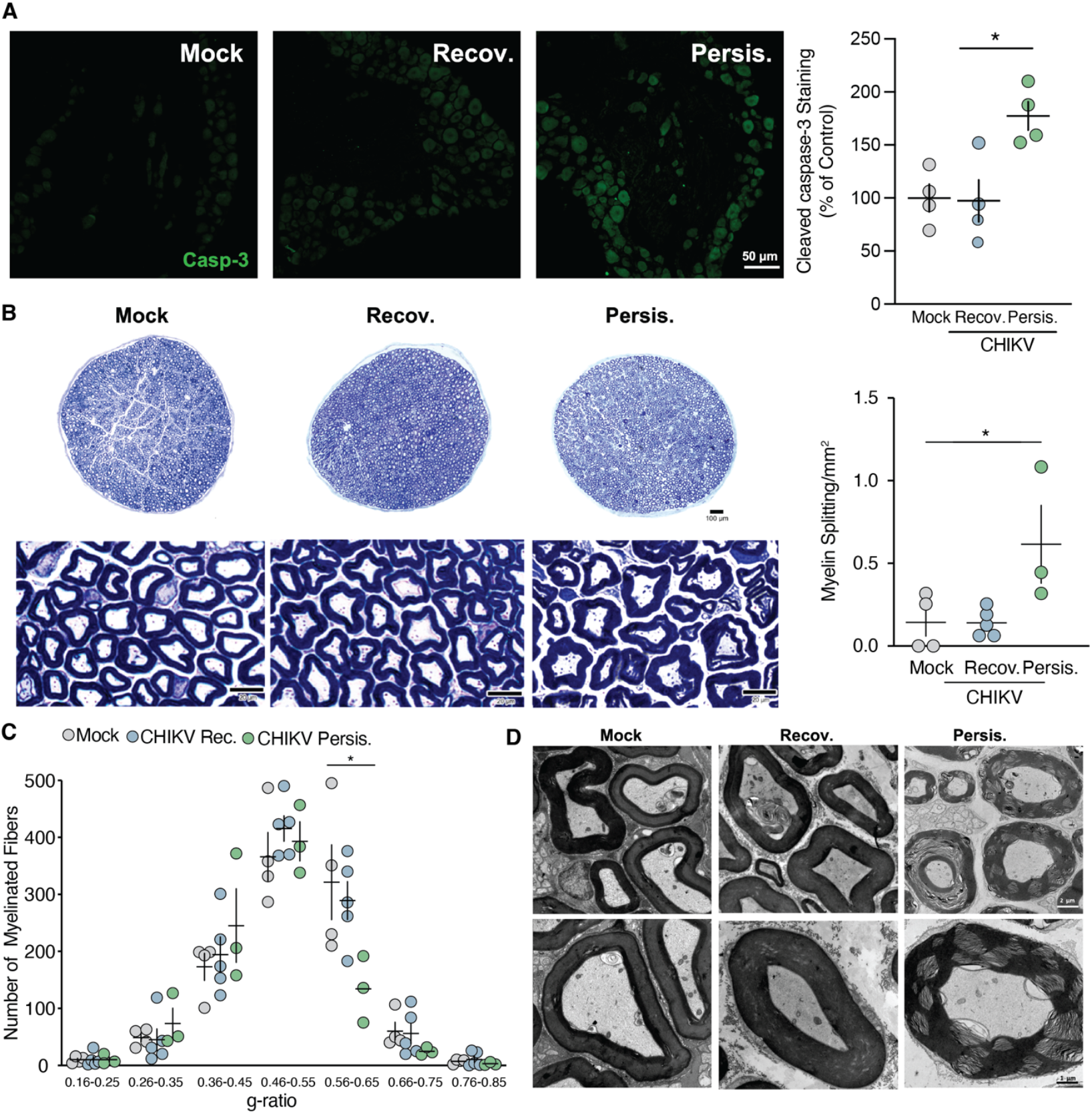
Persistent pain after CHIKV infection is associated with caspase-3 activation in dorsal root ganglia and neuronal stress and peripheral myelin abnormalities. **a**, Cleaved caspase-3 immunostaining in lumbar DRG at 77 dpi from mock, CHIKV-Recov., and CHIKV-Persis. mice, with quantification. **b**, Semithin toluidine blue–stained transverse sections of sciatic nerves at 77 dpi. Quantification of myelin splitting events. **c**, G-ratio distribution of myelinated fibers. **d**, Transmission electron microscopy of sciatic nerves at 77 dpi. Data are the mean and standard error. In **a**, 4 mice per group, Kruskal-Wallis test; **b** (myelin splitting3–5 mice per group, Kruskal-Wallis test; **c** (g-ratio distribution), 4–5 mice per group, two-way ANOVA followed by Tukey’s multiple-comparisons test. *P < 0.05.

Ultrastructural analysis by transmission electron microscopy (TEM) further revealed marked myelin abnormalities in sciatic nerves from the persistent pain mouse group. Myelinated fibers from mock-infected controls and recovered animals presented compact, concentrically arranged myelin lamellae with uniform periodicity and preserved axonal morphology; however, fibers from the persistent group showed pronounced myelin pathology. This included extensive lamellar separation and decompaction, intramyelinic vacuolization, accumulation of electron-dense fibrillar material within split lamellae, and irregular myelin profiles with focal myelin unravelling (**Fig. 5c**). These findings demonstrate that persistent pain following CHIKV infection is accompanied by sustained caspase-3 activation in sensory neurons and structural myelin abnormalities in the sciatic nerve, revealing a chronic peripheral neuropathic phenotype in the absence of gross fiber loss.

### Elevated IFN-γ levels during acute CHIKV infection are associated with chronic pain outcomes in patients

We next examined whether systemic IFN-γ responses during acute CHIKV infection are associated with persistent pain outcomes in patients. To this end, we measured serum levels of TNF-α, IL-6, IL-1β, and IFN-γ in a prospective cohort of individuals who either recovered (n = 20) or developed chronic arthralgia (n = 22) three months post-infection. Odds-ratio analysis revealed that none of the acute-phase clinical symptoms were significantly associated with chronic outcome (**Fig. 6a**). In contrast, we found that IFN-γ levels measured during the acute phase were significantly higher in patients who subsequently developed chronic pain compared with those who recovered. Conversely, concentrations of TNF-α, IL-6, and IL-1β did not differ significantly between groups (**Fig. 6b**). Receiver operating characteristic (ROC) analysis further demonstrated that IFN-γ displayed superior discriminative performance for chronic arthralgia compared with TNF-α, IL-6, and IL-1β (**Extended Data Fig. 5e**). Together, these results indicate that elevated IFN-γ during acute CHIKV infection is a shared feature of both experimental and clinical disease and is associated with an increased risk of persistent pain.

**Fig. 5.**
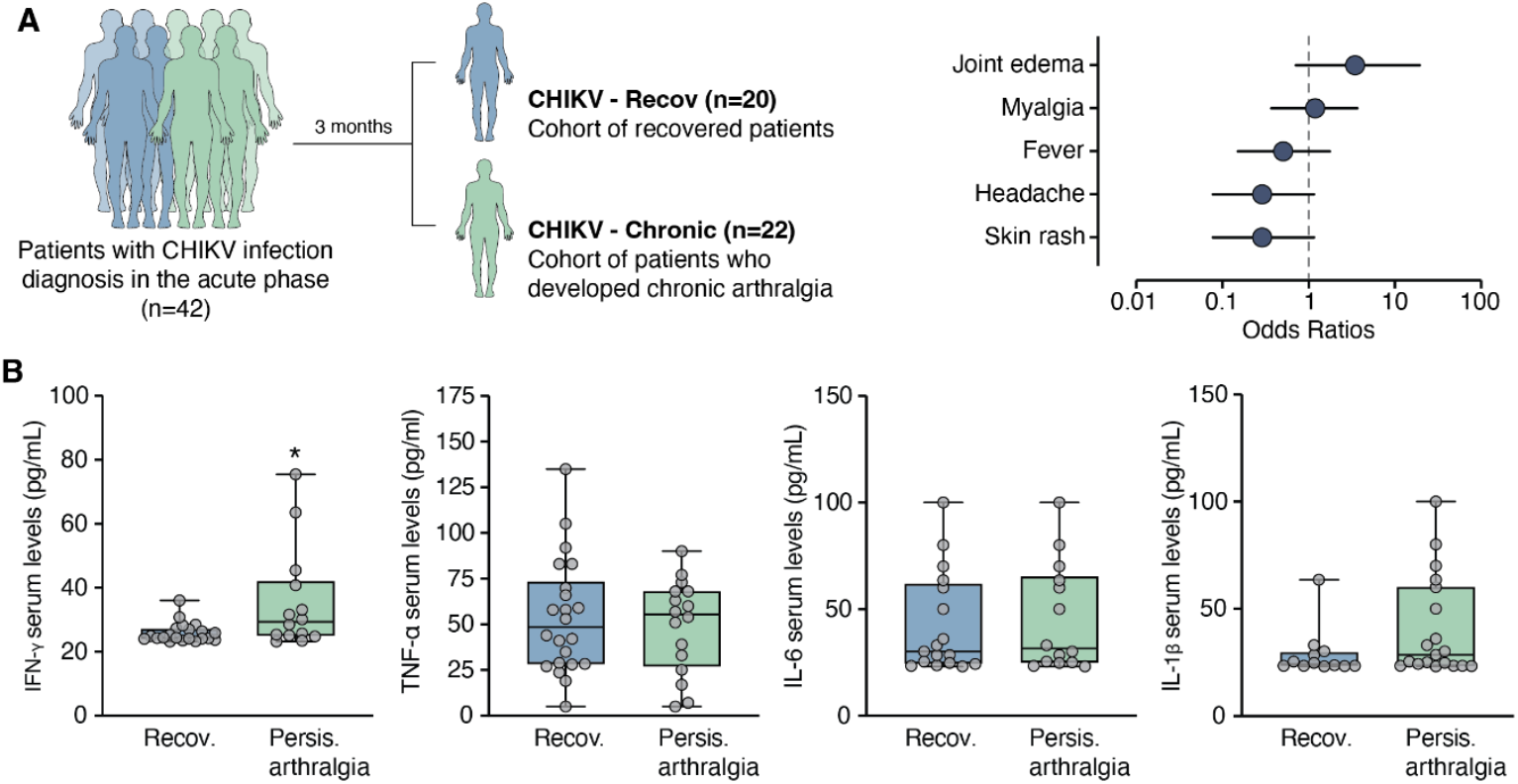
Elevated acute IFN-γ levels are associated with chronic arthralgia after CHIKV infection in humans. **a**, Study design and patient stratification. Individuals with laboratory-confirmed CHIKV infection were enrolled during the acute phase and reassessed 3 months later as recovered (CHIKV-Recov., n = 20) or chronic arthralgia (CHIKV-Chronic, n = 22). Odds ratios for acute-phase clinical symptoms are shown. **b**, Serum cytokine levels measured during the acute phase in CHIKV-Recov. and CHIKV-Chronic patients (TNF-α, IL-6, IL-1β, and IFN-γ). Data are the mean and standard error. In **b**, 14–22 patients per group, a two-tailed unpaired t-test. *P < 0.05.

## Discussion

In this study, we identify IFN-γ as a determinant of chronic pain following CHIKV infection based on murine models and clinical data. Our findings demonstrate that higher levels of IFN-γ during the acute phase correlate with persistent arthralgia. Mechanistically, we show that IFN-γ induces sensory neuron stress and mechanical hypersensitivity independent of viral persistence. This aligns with the clinical absence of detectable viral proteins or RNA in the synovial fluid of patients with chronic joint pain^25^. We also identified CD8^+^ T cells as the predominant source of IFN-γ during the acute phase, consistent with evidence that CHIKV triggers robust CD8^+^ T cell activation in patients^26,27^. Although CD8^+^ T cells have been proposed to clear CHIKV-infected cells, prior studies indicate that their antiviral effects against CHIKV can occur independently of IFN-γ^28^. Collectively, our results establish a causal link between elevated IFN-γ during the acute phase and long-term chronic pain. However, further research is required to distinguish the specific signaling patterns that drive IFN-γ production during early infection and their subsequent impact on nociceptive pathways.

Our findings reveal that persistent pain during the chronic phase of CHIKV infection is likely driven by IFN-γ-mediated injury to peripheral sensory neurons, which disrupts nerve integrity and promotes a neuropathic state. This hypothesis is supported by previous studies demonstrating that IFN-γ acts on dorsal root ganglia and drives neuropathic pain pathways^23,29-31^. Importantly, although CHIKV infection can cause bone erosion and cartilage degeneration^32-34^, we did not observe a correlation between the severity of cartilage degeneration and persistent pain. Likewise, viral RNA levels in joint tissue and dorsal root ganglia were comparable between persistent and recovered animals, indicating that viral persistence does not appear to be the mechanism that leads to chronic pain. In contrast to IFN-γ, acute-phase levels of TNF-α, IL-1β, and IL-6 did not predict pain during the chronic phase. Thus, future population-level studies during CHIKV outbreaks might consider evaluating IFN-γ as a prognostic biomarker for the chronic phase. Furthermore, previous research has shown that higher levels of IL-17 correlate with persistent arthralgia, suggesting that IL-17-driven pathways may serve as an alternative or complementary mechanism alongside IFN-γ-mediated effects^16^.

This study has several limitations. First, our mouse model uses intra-articular CHIKV infection to reproduce key features of CHIKV-induced pain and enables mechanistic interrogation, but we acknowledge that this model may not fully recapitulate the systemic and genetic heterogeneity of chikungunya disease in humans. Second, our analyses primarily focused on peripheral sensory neuron injury; further studies evaluating the potential contributions of central sensitization or spinal circuit remodeling are required. Third, our clinical data on higher IFN-γ levels during the acute phase as a predictor of long-term pain were limited to a single outbreak. Fourth, both our experimental model and patient cohorts were limited to the CHIKV East/Central/South African (ECSA) lineage; further studies utilizing other CHIKV lineages are required to confirm these results. Finally, the mechanism by which downstream intracellular pathways link IFN-γ signaling to sustained neuronal dysfunction that drives pain during the chronic phase remains to be defined.

In conclusion, our findings identify acute IFN-γ-driven neuroimmune signaling as a mechanism linking CHIKV infection to pain in the chronic phase. Moreover, this work expands our knowledge of chikungunya pathogenesis, which may inform future studies focused on identifying therapeutic targets to prevent pain and establishing prognostic biomarkers for the chronic phase.

## Acknowledgments

This work was supported by the Fundação de Amparo à Pesquisa do Estado do Rio de Janeiro (FAPERJ) (E-26/210.779/2021, E-26/210.075/2023, E-26/200.291/2025, E-26/2020.752/2019, E-26/010.000169/2020, SEI-260003/001141/2020, E-26/010.002260/2019), the Coordenação de Aperfeiçoamento de Pessoal de Nível Superior (CAPES) (88887.657702/2021-00), the Conselho Nacional de Desenvolvimento Científico e Tecnológico (CNPq) (312891/2018), the CAPES/STINT Brazil–Sweden Cooperation Program (88887.465506/2019-00), and the Fundação de Amparo à Pesquisa do Estado de São Paulo (FAPESP) (2024/22135-0). WMdS was supported by Wellcome Trust – Digital Technology Development Award in Climate Sensitive Infectious Disease Modelling (#226075/Z/22/Z). We thank João Batista Calixto and Marcelo Alves Mori for critical reading of the manuscript, and Ana Claudia Rangel, José Nilson dos Santos, Daiane Torres, and Bianca de Lima Almeida for technical support, including assistance with primary cell cultures. We thank Jacqueline A. Tida (www.plotmyscience.com) for figure editing.

## METHODS

### Animals

Adult male and female Swiss mice were used in most experiments. In some experiments, C57BL/6J wild-type (WT) and *Ifng*^*-/-*^ mice^35^ were used. All mice were 8–12 weeks old. Animals were housed in groups of five per cage with free access to food and water, under 12 hours light/dark cycle, with controlled temperature and humidity. All procedures followed were approved by the Institutional Animal Care and Use Committee of the Federal University of Rio de Janeiro, Brazil (Approval: A19/24-008-21).

### Viruses

CHIKV strain BHI3768 was isolated from a febrile Brazilian patient during the 2016 outbreak, and viral stocks were generated as previously described^36^. To generate CHIKV stock, C6/36 cells were infected at a multiplicity of infection (MOI) of 0.01. After 30 hours of infection, the culture supernatant was then collected and clarified by centrifugation at 500 g for 10 min. Clarified supernatants were aliquoted and stored at −80°C. Viral titers of the resulting stocks were determined by plaque assay in Vero cells.

### CHIKV inoculation

Mice were anesthetized with isoflurane and inoculated intra-articular in the left knee joint with 10 µL of CHIKV strain BHI3768 at 10^6^ PFU. Control animals received an equivalent volume of virus-free C6/36 medium (mock control).

### RNA extraction and qPCR

Total RNA was extracted from knee joints and DRG using TRIzol Reagent (Cat No. 15596026, Invitrogen, USA), according to the manufacturer’s instructions. RNA concentration and purity were assessed using a Thermo Scientific NanoDrop spectrophotometer by measuring absorbance ratios at 260/280 and 260/230 nm. RNA extractions with absorbance ratios ≥1.8 were used. 1µg of isolated RNA was reverse-transcribed using the High-Capacity cDNA Reverse Transcription Kit (Cat No. 4368814, Thermo Fisher Scientific, USA). Quantitative PCR was performed using Power SYBR Green PCR Master Mix (Cat No. 4368577, Life Technologies, USA) on a QuantStudio 3 Real-Time PCR System (Thermo Fisher Scientific). Gene expression was quantified using the ΔΔCt method and normalized to β-Actin. Target genes included inflammatory and matrix remodeling markers (*Tnf, Il6, Ifng, Ifna, Ifnb, Mmp13, Adamts4, Rankl, Opg)* and neuronal stress markers such as *Atf3*. Viral load detection was performed using the TaqMan Mix kit (Cat No. 4444556, Thermo Fisher Scientific, USA) following the manufacturer’s protocol, employing primers and probe specific for the CHIKV nsp1 gene (**Extended Table 1**). Cycle threshold values were used to calculate the equivalent PFU/μg of total RNA after conversion, using a standard curve with serial 10-fold dilutions of a CHIKV stock. Primer sequences for all targets are listed in **Extended Table 2**.

### Viral plaque assay

Viral titers were quantified by plaque formation assay. Tissues were weighed and homogenized in Dulbecco’s Modified Eagle’s Medium (DMEM) supplemented with penicillin and streptomycin at a ratio of 0.2 mg tissue/µL sample. After homogenization, 25 µL of each sample was serially diluted in serum-free DMEM High Glucose. Semi-confluent Vero cells seeded in 24-well plates were inoculated with 200 µL of each dilution (10^−1^ to 10^−6^) and incubated for 1 hour at 37°C with 5% CO_2_. Following adsorption, wells were overlaid with 1 mL of DMEM containing 1% carboxymethylcellulose, 1% Fetal Bovine Serum (FBS), and 1% penicillin/streptomycin and incubated for 40 hours under standard culture conditions. Cells were then fixed with 10% formaldehyde for 1 hour, washed, and stained with 1% crystal violet in 20% ethanol for 20 min. Plaques were counted, and titers expressed as plaque-forming units (PFU).

### Immunohistochemistry for dsRNA

Knee joints were fixed in 4% paraformaldehyde, decalcified in ethylenediaminetetraacetic acid (EDTA), embedded in paraffin, and sectioned sagittally at 5 µm. Sections were deparaffinized, rehydrated, and subjected to antigen retrieval in citrate buffer (pH 6.0) for 30 minutes at 95–98°C. Tissue permeabilization was performed with 0.5% Triton X-100 for 30 minutes, followed by blocking in 0.2% Triton X-100, 3% bovine serum albumin (BSA), and 5% normal goat serum (NGS) for 2 hours at room temperature. Primary antibodies against dsRNA (clone J2; Cat no. Netherlands, Nordic-MUbio, Netherlands) were diluted in primary antibody diluent and applied overnight at 4°C. After washing, sections were incubated with Alexa Fluor™ 488 goat anti-mouse IgG secondary antibody (Cat No. A28175 Invitrogen, USA) for 1 hour at room temperature. Nuclei were counterstained with 4’,6-diamidino-2-phenylindole (DAPI), and slides were mounted with fluorescence mounting medium for imaging.

### Pain sensitivity assessment

Mechanical sensitivity was assessed using calibrated von Frey filaments (0.07–2.0 g) applied to the plantar surface of the ipsilateral hind paw^37^. Paw withdrawal thresholds were determined using the up-and-down method^38^. Cold sensitivity was evaluated by applying a drop of acetone (approximately 50 µL) to the plantar surface, and the duration of nocifensive behaviours (shaking, licking, or lifting of the paw) was recorded over 60 seconds^39^. Mice were habituated to the testing environment for at least 1 hour before each session. All evaluations were performed at baseline and at defined time points post-infection by experimenters blinded to experimental groups.

### Gabapentin treatment and mechanical sensitivity assessment

Animals infected intra-articularly with 10^6^ PFU/mL of CHIKV were longitudinally evaluated for mechanical sensitivity using the von Frey test. At 28 days post-infection, when mechanical hypersensitivity was fully established, animals were treated with gabapentin (100 mg/kg) administered orally. Control animals received the drug vehicle, 0.5% carboxymethylcellulose (CMC), also administered orally. Mechanical sensitivity was assessed using von Frey filaments at 1, 3, 5, and 24 hours following drug or vehicle administration.

### Histology and cartilage damage scoring

Knee joints were harvested, fixed in 4% paraformaldehyde, decalcified in EDTA, and embedded in paraffin. Serial sagittal sections (5 µm) were stained with hematoxylin and eosin (H&E), safranin-O/fast green, or toluidine blue using standard protocols. Cartilage pathology in H&E-stained sections was scored by two independent blinded observers, as previously described^40^. For safranin-O/fast green and toluidine blue staining, cartilage matrix content was quantified using ImageJ software by measuring the stained area and intensity relative to total cartilage area in defined regions of interest (ROIs).

### Flow cytometry

Mock (uninfected) and CHIKV-infected mice were anesthetized with ketamine/xylazine and subjected to two consecutive intra-articular lavages of the knee using 10 µL of phosphate-buffered saline (PBS) supplemented with 5% FBS. Synovial fluid samples were collected and stained with fluorochrome-conjugated monoclonal antibodies for immunophenotyping of leukocyte populations. The following antibodies were used: anti-CD4 PE (Cat No. 12-0049-42, eBioscience, USA), anti-CD8 APC (Cat No. 17-0084-82, eBioscience, USA), anti-CD11b FITC (Cat No. 11-0118-42, eBioscience, USA), and anti-CD45 PE-Cy5 (Cat No. 35-0451-82, eBioscience, USA). Samples were incubated with the antibody mixture for 30 minutes at room temperature in the dark, washed with PBS containing 5% FBS, and resuspended in staining buffer. Flow cytometric acquisition was performed using a FACSCanto™ cytometer (BD Biosciences), and data were analyzed with FlowJo™ software (BD Biosciences). Compensation was performed using single-stained controls for each fluorochrome. Cells were first gated based on forward and side scatter (FSC-A versus SSC-A) to exclude debris. Doublets were excluded using FSC-A versus FSC-H. The main leukocyte population was subsequently identified based on FSC and SSC characteristics. CD45^+^ leukocytes were then gated and further subdivided into CD4^+^ T cells (CD45^+^CD4^+^CD8^−^), CD8^+^ T cells (CD45^+^CD8^+^CD4^−^), and phagocytic cells (CD45^+^CD11b^+^).

### Intracellular IFN-γ detection by flow cytometry

Mock- and infected mice were anesthetized with ketamine/xylazine and subjected to two consecutive intra-articular lavages of the knee using 10 µL of PBS supplemented with 5% FBS. Synovial fluid cells were collected and stimulated with Cell Stimulation Cocktail (Cat No. 00-4970-03, eBioscience, USA) for 3 hours in the presence of brefeldin A to inhibit cytokine secretion. Following stimulation, cells were stained for surface markers using anti-CD4 PE (Cat No. 12-0049-42, eBioscience, USA), anti-CD8 eFluor450 (Cat No. 48-0081-82, eBioscience, USA), anti-CD45 PE-Cy5 (Cat No. 35-0451-82, eBioscience, USA), and anti-CD49b FITC (Cat No. 11-0491-82, eBioscience, USA). Cells were then fixed and permeabilized using the BD Cytofix/Cytoperm™ kit (Cat no., 554714, BD Biosciences, USA) according to the manufacturer’s instructions and stained intracellularly with anti–IFN-γ APC (Cat No. 17-7311-82, eBioscience, USA). Samples were washed, resuspended in PBS, and acquired on a BD FACSCanto™ flow cytometer. Data were analyzed using FlowJo software (BD Biosciences). Compensation was performed using single-stained controls. Cells were first gated based on forward and side scatter (FSC-A versus SSC-A) to exclude debris, followed by exclusion of doublets using FSC-A versus FSC-H. CD45^+^ leukocytes were subsequently gated and subdivided into CD4^+^ T cells, CD8^+^ T cells, and CD49b^+^ cells. Intracellular IFN-γ expression was then quantified within each gated population.

### Gating strategy

Doublets were excluded by FSC-A × FSC-H. Live cells were selected based on FSC × SSC, followed by gating on CD45^+^ leukocytes. These were subdivided into CD4^+^, CD8^+^, and CD49b^+^ populations, and IFN-γ expression was quantified within each subset.

### Intra-articular injection of recombinant IFN-γ

Recombinant murine IFN-γ (Cat No. 485-MI/CF, R&D Systems, USA) was administered intra-articular into the left knee joint of naive mice at a dose of 300 U/site every 48 hours over a 7-day period. This dosing regimen was selected based on preliminary titration experiments showing that a higher dose (600 U/site) did not produce additional effects compared with the 300 U dose, and was consistent with intra-articular doses previously used for other recombinant cytokines in the literature^41,42^. Mechanical sensitivity was assessed using calibrated von Frey filaments, as described above.

### DRG neuron culture and stimulation

DRG neurons were isolated from adult mice and cultured for 24 hours following a previously described protocol^43^. Cells were then stimulated with recombinant IFN-γ (1 ng/mL or 8,4 U/mL; Cat No. #485-MI/CF, R&D Systems, USA) or capsaicin (1 µM, positive control) for 60 minutes. Neurons were fixed and immunostained for p-ERK (Anti-p-ERK, Cat No. 4370S, Cell Signaling, USA), ATF3 (Anti-ATF3, Cat No. NBP1-02935, NovusBio, USA), and NeuN (Anti-NeuN, Cat No. 24307S, Cell Signaling, USA). Confocal images were acquired to quantify the percentage of marker-positive neurons. In parallel, *Atf3* mRNA expression was assessed by RT-qPCR.

### IFN-γ receptor blockade

Pharmacological inhibition of IFN-γ signaling was achieved via intra-articular administration of a monoclonal anti-IFN-γ receptor antibody (anti-CD119; clone GR-20, Cat No. 16-1193-85, Invitrogen, USA). The antibody was injected at 0.5 and 5 µg/site^44,45^ into the left knee joint at the time of infection and subsequently every 48 hours for a total of 4 doses. Control animals received an equivalent amount of isotype-matched IgG (Cat No. 16-4321-82, Invitrogen, USA). In parallel, Ifng^-/-^ mice^35^ were used to genetically validate the role of IFN-γ signaling in CHIKV-induced responses.

### IFN-γ quantification

Systemic levels of IFN-γ were measured in mouse serum samples collected at defined time points using a commercially available ELISA kit (Cat No. DY485-05, R&D Systems, USA), following the manufacturer’s instructions. Samples were run in triplicate, and cytokine concentrations were calculated based on standard curves generated in each plate.

### Semithin section preparation and morphometric analysis

Semithin nerve analyses were performed as previously described^46^. Transverse semithin sections (300–500 nm) were collected on glass slides, stained with toluidine blue, and imaged at 100× magnification using a light microscope equipped with a digital camera. Qualitative and quantitative assessments of myelinated fibers were performed along the length of each nerve. For morphometric analysis, five fields per nerve were acquired at 100× magnification to quantify myelin abnormalities and G-ratio. Additionally, panoramic images were analyzed from 40× micrographs of each nerve, and the total number of myelinated fibers was quantified.

### NF200 Immunostaining of the sciatic nerve

Dissected sciatic nerves were fixed in 4% paraformaldehyde for 24 hours, washed in PBS (pH 7.4) for 24 hours, and cryoprotected in 10%, 20%, and 30% sucrose solutions (48–72 hours each). Tissues were embedded, flash-frozen in liquid nitrogen, and sectioned transversely at 12 µm using a cryostat (Leica CM1850). Sections were washed twice in 0.3% PBST and incubated in a blocking solution (PBS containing 0.3% Triton X-100 and 10% normal goat serum) for 1 hour at room temperature. Samples were then incubated with the primary antibody anti-NF200 (1:1000; Cat No. N4142, Sigma-Aldrich, USA) for 24 hours at 4°C. After PBS washes, sections were incubated with Alexa Fluor™ 488 goat anti-mouse IgG (1:500; Cat No. A28175, Molecular Probes, USA) for 2 hours at room temperature, washed, and counterstained with DAPI (Cat No. D9542, Sigma-Aldrich, USA). Slides were mounted with Fluoromount (Cat No. 17984-25, Electron Microscopy Sciences, USA) and stored at ™20°C. Omission of the primary antibody served as a negative control. Fluorescence images were acquired on a Zeiss microscope using a 20× objective. The images were processed and analyzed using Fiji (ImageJ). Mean fluorescence intensity was measured by selecting regions of interest (ROIs) and using the Analyze > Measure function to record the Mean Gray Value of NF200 immunoreactivity.

### Cleaved caspase-3 immunostaining of the DRG

Dissected DRGs were fixed in 4% paraformaldehyde for 24 hours, washed in PBS (pH 7.4) for 24 hours, and cryoprotected in 10%, 20%, and 30% sucrose solutions (48–72 hours each). Tissues were embedded, flash-frozen in liquid nitrogen, and sectioned transversely at 12 µm using a cryostat (Leica CM1850). Sections were washed twice in 0.3% PBST and incubated in a blocking solution for 1 hour at room temperature. Samples were then incubated with the primary antibody anti-cleaved caspase-3 (1:300; Cat No. 9661S, Cell Signaling, USA) for 24 hours at 4 °C. After PBS washes, sections were incubated with Alexa FluorTM 488 goat anti-mouse IgG (1:500; Cat No. A28175, Molecular Probes, USA) for 2 hours at room temperature, washed, and counterstained with DAPI (Cat No. D9542, Sigma-Aldrich, USA). Slides were mounted with Fluoromount (Cat No. 17984-25, Electron Microscopy Sciences, USA) and stored at ™20°C. Omission of the primary antibody served as a negative control. Fluorescence images were acquired on a Zeiss microscope using a 20× objective. The images were processed and analyzed using Fiji (ImageJ). Mean fluorescence intensity was measured by selecting ROIs and using the Analyze > Measure function to record the Mean Gray Value of cleaved caspase-3 immunoreactivity.

### Electron microscopy

Sciatic nerves were fixed in 2.5% glutaraldehyde diluted in 0.1 M cacodylate buffer for 24 hours, washed, and post-fixed for 2 hours in 1% osmium tetroxide containing 0.8% potassium ferrocyanide in 0.1 M cacodylate buffer (pH 7.4). Following, samples were washed and dehydrated in an acetone graded series before being embedded in an increasing concentration of Spurr. Spurr-embedded samples were hardened at 60°C for 72 hours, and 80-nm ultrathin sections were prepared and attached to copper grids. Lead citrate and uranyl acetate were used for post-staining, and samples were observed with a transmission electron microscope (HITACHI HT 7800).

### Patients with CHIKV infection and cytokine analysis

A retrospective cohort of patients with acute CHIKV infection (i.e., ≤ 14 days post onset symtomys) during the outbreak in Rio de Janeiro City, Brazil (2018–2019) was analyzed. Forty-two participants over 18 years (median: 35 years old; interquartile: 49-66), sex-ratio of 0.6 female-to-male. All patients with laboratory-confirmed CHIKV infection were diagnosed by RT-qPCR or IgM ELISA, plus clinical manifestations such as fever, musculoskeletal manifestations, rash myalgia. Individuals with a history of chronic pain were excluded. Participants were followed up using a structured questionnaire to assess the incidence of chronic arthralgia (joint pain lasting ≥3 months after diagnosis) and its association with joint pain characteristics during the acute phase, as well as levels of inflammatory markers TNF-α, IL-6, IL-1β, and IFN-γ in serum samples were quantified using Human IL-1β/IL-1F2 Quantikine ELISA Kit (Cat No. DLB50, R&D Systems, USA) and Milliplex® MAP Kit High Sensitivity Magnetic Bead Cytokine Panel for TNF-α and IL-6 (Cat No. HCYTA-60K, Merck, USA), both according to the manufacturer’s instructions. All procedures were approved by the Ethics committee of the Federal University of Rio de Janeiro (Approval: 42340920.7.0000.5259), and written informed consent was obtained from all participants.

### Statistical analysis

Data are presented as mean and standard error unless otherwise indicated. Group comparisons were performed using two-tailed unpaired t-tests or one- or two-way ANOVA followed by appropriate post hoc tests, as specified in the figure legends. When the data did not meet the assumptions of normality, nonparametric tests were used. Normality was assessed using Shapiro–Wilk tests. Statistical significance was set at P < 0.05. For viral load kinetics, independent cohorts of mice were euthanized at each time point for tissue collection and analysis. For some qPCR targets, sample sizes varied slightly due to limited cDNA availability per sample. No biological samples were excluded from analysis except when predefined technical quality criteria were not met. Statistical analyses were conducted using GraphPad Prism version 10 (GraphPad Software, USA) and R version 4.4.2 (R Foundation for Statistical Computing, Austria).

