## Supplementary material for "Acute IFN-γ responses drive sensory neuron injury and chronic pain after chikungunya virus infection": Figure S (from another version of this preprint: version 1, 2026-02-03)

### Supplementary Figures Legends:

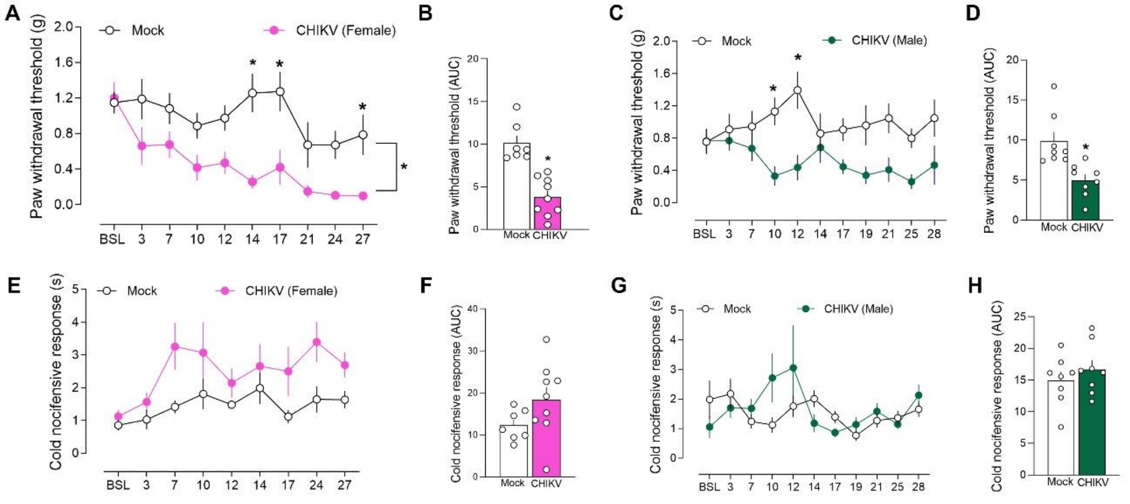


Suppl. Fig 1

**Figure S1. CHIKV infection induces mechanical and cold allodynia in male and female mice. (A–D)** Time course of paw withdrawal thresholds (PWT, g) to von Frey stimulation in Mock- and CHIKV-inoculated mice (0-28 dpi) **(A)** Female mice; **(B)** area under the curve (AUC) corresponding to panel A; **(C)** male mice; and **(D)** AUC corresponding to panel C. CHIKV infection significantly reduced PWT compared with Mock at multiple time points in both sexes. Data are presented as mean ± SEM. Two-way ANOVA with Sidak's multiple-comparisons test, $P < 0.05$. $n = 7–9$ animals per group.

**(E–H)** Time course of cold nocifensive responses (s) to acetone stimulation in Mock- and CHIKV-inoculated mice (0-28 dpi). **(E)** Female mice; **(F)** AUC corresponding to panel E; **(G)** male mice; and **(H)** AUC corresponding to panel G. Only CHIKV-infected female mice exhibited increased cold nocifensive responses. Data are presented as mean ± SEM. Two-way ANOVA with Sidak's multiple-comparisons test in (E) and (G), n.s. Two-tailed unpaired *t* test in **(F)**, $P = 0.0009$, and **(H)**, n.s. $n = 7–9$ animals per group.

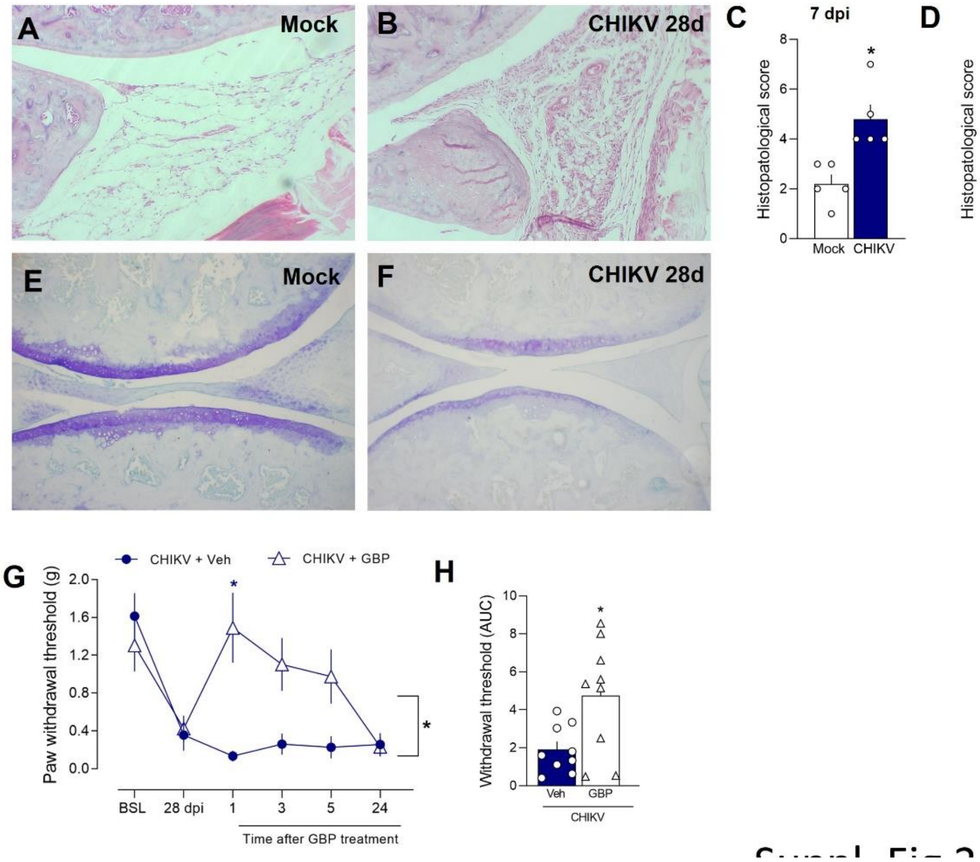


**Figure S2. Immune cell infiltration, cartilage integrity, and pharmacological modulation of mechanical allodynia after CHIKV infection**

**(A–D)** Immune cell infiltration in the infected joint. Representative images and quantification of immune cell infiltrates in the articular region at 7 and 28 dpi show robust inflammation at 7 dpi **(C)**, with persistent immune cell presence at 28 dpi **(D)**. Two-tailed unpaired *t* test in **(C)** $P = 0.005$ and **(D)** $P = 0.028$. $n$ = 5-8 animals per group.

**(E, F)** Toluidine blue staining of knee joint sections reveals cartilage alterations following CHIKV infection. Representative images and quantitative analysis (see Figure 1O) indicate increased cartilage damage in infected animals compared with Mock controls.

**(G, H)** Effect of gabapentin treatment on CHIKV-induced mechanical allodynia at 28 dpi. **(G)** Time course of paw withdrawal thresholds (PWT, g) to von Frey stimulation in CHIKV-infected mice following gabapentin administration (p.o., 100 mg/kg). **(H)** Area under the curve (AUC) analysis. A single dose of gabapentin attenuates mechanical allodynia in CHIKV-infected mice. Two-way ANOVA with Sidak's multiple-comparisons test in **(G)**, $P < 0.05$; two-tailed unpaired *t* test in **(H),** $P = 0.017$. $n$ = 9 animals per group.

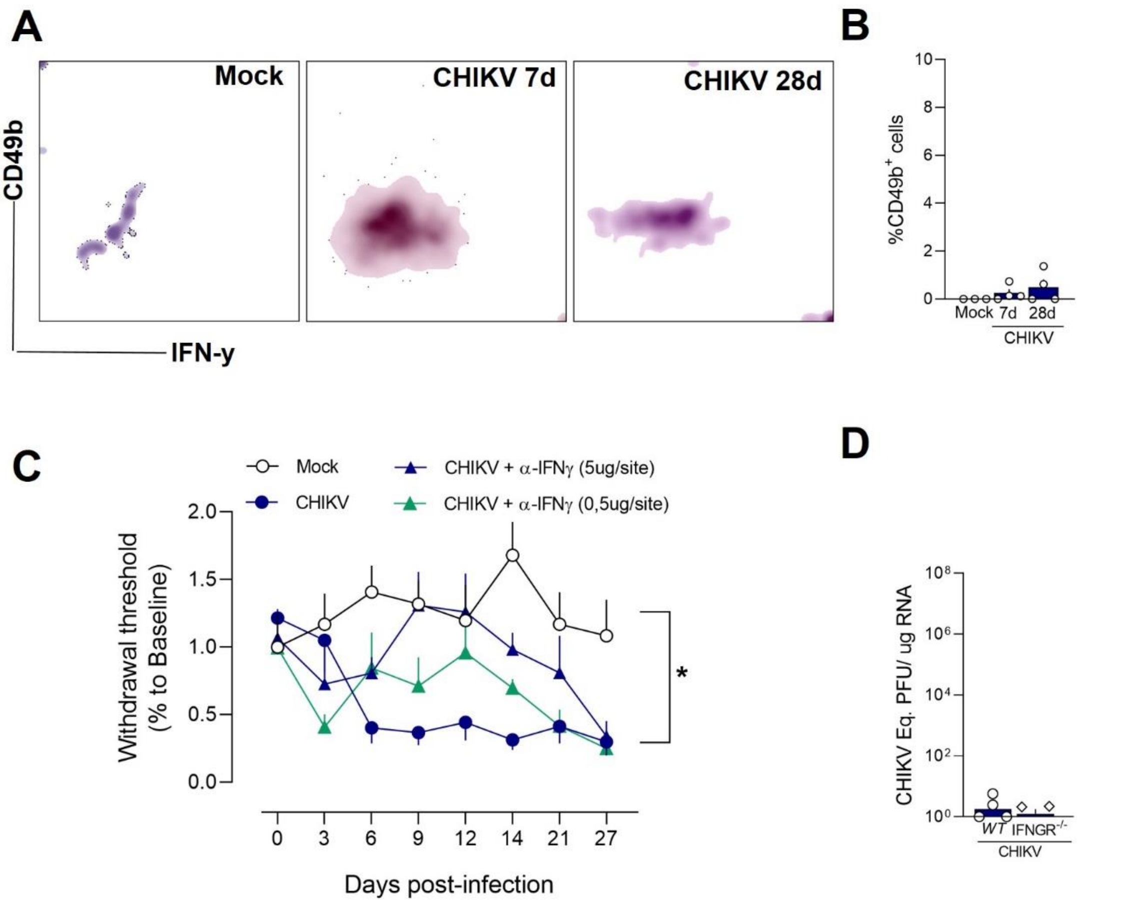


Suppl. Fig 3

**Figure S3.NK cell–derived IFN-γ, IFN-γ receptor blockade, and viral load in DRG after CHIKV infection**

**(A, B)** Flow cytometry analysis of IFN-γ–producing NK cells in the synovial fluid of the infected knee joint at 7 and 28 dpi. **(A)** Representative gating strategy and plots; **(B)** quantification of IFN-γ–secreting NK cells. One-way ANOVA with Tukey's multiple-comparisons test; n.s. n = 4 animals per group.

**(C)** Effect of intra-articular blockade of IFN-γ signaling on CHIKV-induced mechanical allodynia. Paw withdrawal thresholds (PTW, g) measured by von Frey testing show that intra-articular administration of an anti-CD119 antibody (5 µg/site or 0.5 µg/site) attenuates mechanical allodynia, with the 5 µg dose showing greater efficacy. Two-way ANOVA with Tukey's multiple-comparisons test; $P < 0.05$. n = 8–9 animals per group.

**(D)** CHIKV RNA levels in dorsal root ganglia (DRG) from IFN-γ–deficient (*Ifng*$^{-}$/$^{-}$) mice. No differences in viral load were detected between groups, indicating that IFN-γ signaling does not influence viral persistence in the DRG. Two-tailed unpaired *t* test; n.s. n = 4–6 animals per group.

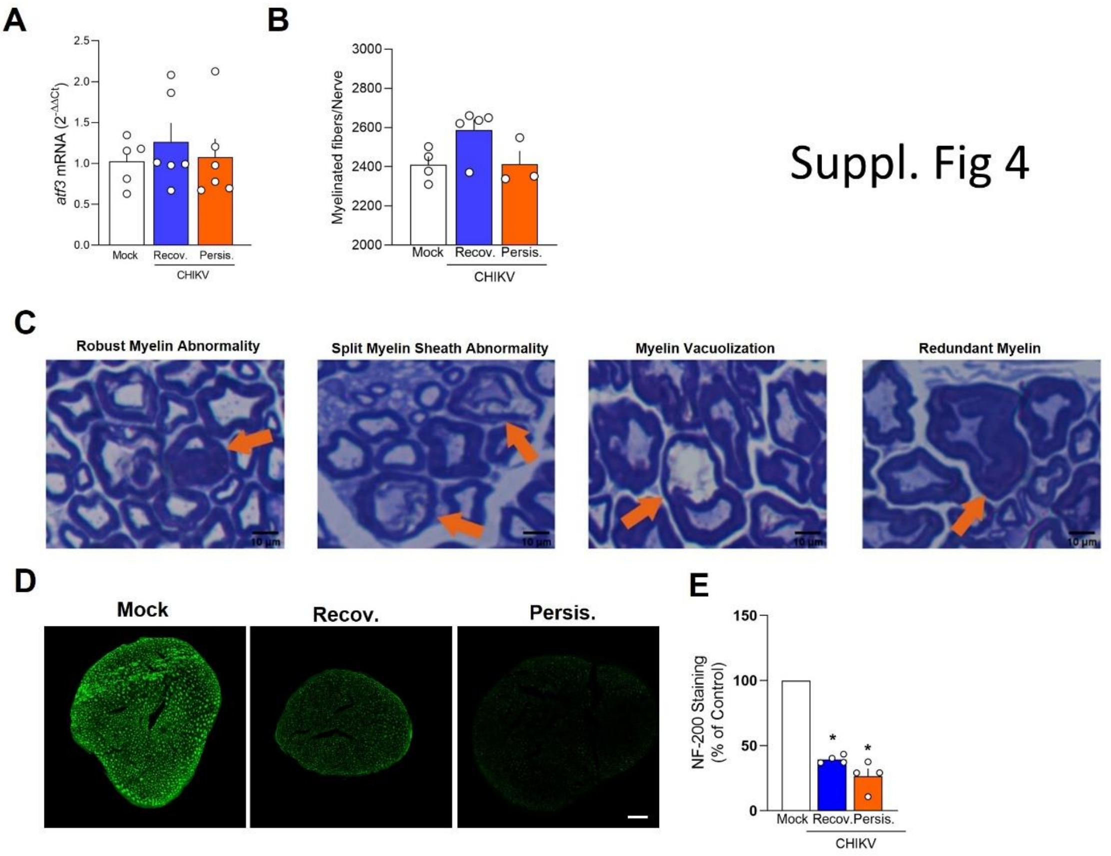


**Figure S4. Neuronal stress markers and sciatic nerve structural alterations associated with persistent mechanical allodynia after CHIKV infection**

**(A)** qPCR analysis of *Atf3* mRNA expression in dorsal root ganglia (DRG) from Mock, recovered (CHIKV-Recov.), and persistent (CHIKV-Persis.) mice. No significant differences were observed among groups. One-way ANOVA with Tukey's multiple-comparisons test, n.s. n = 5-6 animals per group.

**(B)** Total number of myelinated fibers in the sciatic nerve from Mock, CHIKV-Recov., and CHIKV-Persis. mice. Quantitative analysis revealed no differences in overall myelinated fiber density across groups. One-way ANOVA with Tukey's multiple-comparisons test, n.s. n = 4 animals per group.

**(C)** Representative images illustrating myelin abnormalities in the sciatic nerve of CHIKV-Persis. mice, including split myelin sheaths, myelin vacuolization, and redundant myelin, compared with Mock and CHIKV-Recov. groups.

**(D, E)** Neurofilament integrity in the sciatic nerve. **(D)** Representative immunofluorescence images of NF200 and **(E)** corresponding quantification show reduced neurofilament signal in CHIKV-Persis. mice compared with Mock and CHIKV-Recov. groups, consistent with axonal alterations associated with persistent mechanical allodynia. One-way ANOVA with Tukey's post hoc test; P = 0.041. n = 4 animals per group.
